# CRISPR-Cas9-induced genetic mosaicism in three species of the microcrustacean *Daphnia*

**DOI:** 10.64898/2026.01.26.701744

**Authors:** Sen Xu, Swatantra Neupane, Li Wang

## Abstract

Genetic mosaicism can arise from *in vivo* CRISPR-Cas9 gene editing, especially in the embryos. This study evaluates the extent of genetic mosaicism resulted from CRISPR-Cas9-mediated knockout for 11 genes in the freshwater microcrustacean *Daphnia magna, D. pulex*, and *D. sinensis*. Based on extensive genotyping data of the asexually produced progenies of successfully edited females, we find strong evidence of mosaicism in 9 of these genes. The genotyping data also suggest the gene editing activity can take place as early as the one-cell embryo stage and extends into the 32-cell and later stages. This study establishes genetic mosaicism as an important feature of Cas9-mediated gene editing in *Daphnia*.

## Introduction

CRISPR-Cas-mediated genetic modification can lead to mosaicism, i.e., different genotypes exist in the same individual (Davies 2019; Mehravar et al. 2019). This effect is often exacerbated when introducing CRISPR-Cas editing system into embryos, where CRISPR-Cas can continuously edit genes in different cells arising from different stages of embryo development. Although mosaicism is often considered as a drawback for gene editing, mosaicism occurring in primary germline cells can lead to different loss-of-function allelic modifications, which can be useful for establishing multiple mutant allelic genotypes and functional analysis. Furthermore, interpreting mosaicism in the context of primordial germ cell division and embryo development can help us understand the dynamics of gene editing activities in embryos.

CRISPR-Cas9 has been demonstrated to be a powerful tool for generating gene-knockout mutants for the freshwater microcrustacean *Daphnia* (Ismail et al. 2018; Xu et al. 2025). *Daphnia* typically reproduces via cyclical parthenogenesis, i.e., females can reproduce asexually under favorable environments and switch to sexual reproduction in response to adverse environmental conditions (Xu et al. 2025). As an emerging model system of genetics and genomics (Miner et al. 2012; Ebert 2022), CRISPR-Cas9 gene editing has been successfully carried out in *Daphnia* by microinjecting CRISPR-Cas9 plasmid/mRNA/RNP into the asexual, directly developing embryos (Nakanishi et al. 2014; Xu et al. 2025). An intriguing observation arising from previous CRISPR-Cas-mediated gene knock in the species *Daphnia pulex* is that different genotypes appear in the asexual progenies of gene-edited founder females, while in theory asexual progenies of a female *Daphnia* should be genetically identical. For example, when knocking out the gene SCARLET, which is involved in the production of black pigment in the eye, both black-eyed and clear-eyed (i.e., loss of black pigment) offspring are found in the same broods of genetically edited females.

These observations pose some interesting questions. For example, despite our intention that injection of CRISPR-Cas into freshly deposited embryos could lead to editing in the one-cell stage of embryo, the genetic mosaicism in the offspring indicates CRISPR-Cas often takes action much later than the one-cell stage. It would be useful to understand the major time window of CRISPR-Cas editing activities to guide gene editing experiments. Furthermore, it remains to be investigated whether this type of mosaicism is a general feature for gene editing in *Daphnia* or unique to the SCARLET gene in *D. pulex*.

To address these questions, we first introduce the process of differentiation of germ cells during embryo development in *Daphnia*. Once deposited into the brood chamber, the asexual embryos of *Daphnia* are in the metaphase I of a modified, abortive meiosis and completes the abortive meiosis and the first cell division about 40 minutes post ovulation (Ojima 1958; Zaffagnini and Sabelli 1972; Hiruta et al. 2010). From then the embryo goes through a round of cleavage every 30 minutes at 25°C (Sagawa et al. 2005). Based on the study by Sagawa et. al. (2005), the primordial germ cells (PGCs) are specified as early as in the 8-cell stage, and two PGCs appear at the end of the 32-cell stage, then PGCs proliferate into 4 cells at the 64-cell stage, 8 cells at the 128-cell stage, and so on. PGCs eventually settle into regions where ovaries form along both sides of the gut. In adult females, oogonia are at the most posterior end of the ovary (Hiruta et al. 2010; Hiruta and Tochinai 2012), without any ovarioles as found in insects.

Furthermore, we collected intensive genotype data of gene knockout experiments of the SCARLET gene and several other genes in *D. magna, D. pulex*, and *D. sinensis*. The genotype data from the offspring of gene edited females allow us to make meaningful inferences about the timing and action modality of CRISPR-Cas and whether mosaicism is a common feature of CRISPR-Cas9 editing in *Daphnia*.

## Materials and Methods

### Experimental animals

We maintained a healthy culture of adult females of the *Daphnia magna* isolate LRV01 (Denmark) and *D. pulex* isolate Tex23 (Michigan, USA). We kept Tex23 animals in artificial lake water COMBO (Kilham et al. 1998) at 20 °C, whereas LRV01 in 4x concentrated COMBO. The culture of both species was under a 12:12 (light:dark hours) photoperiod. We fed the animals with the green algae *Scenedesmus obliquus* every day and removed the newly born babies every other day to prevent overcrowding to keep female *Daphnia* asexually reproducing.

### CRISPR-Cas9 gene knockout experiments

Our gene knockout experiments include the following genes for *D. magna*, SCARLET, DNMT3A, DNMT3B, CLOCK, PRDM8, DMRT1, and DMRT2 (**Supplementary Table 1**). For *D. pulex*, our target genes include DFH, Insulin-like receptor (**Supplementary Table 1**). For *D. sinensis*, we tested the ISM1 (Isthmin 1) and TM2D2 (TM2 Domain Containing 2) genes. For each gene except CLOCK, we designed two guide RNAs targeting early exons using the guide RNA design tool from Integrated DNA Technologies based on the genomic sequences of LRV01 or Tex23 (**Supplementary Table 1**). CLOCK gene had a single guide RNA. We fused Cas9 nucleases (Integrated DNA Technologies, catalog number 1081059) and chemically synthesized gRNAs (Integrated DNA Technologies) at room temperature. The formed RNP contains each sgRNA at a concentration of 125 ng/μl and Cas9 nuclease at 600 ng/μl. We *in vitro* tested whether the RNP can effectively cut the target site by incubating the RNP with PCR amplicons containing target sites. Through microinjection, we deliver RNP into freshly ovulated asexual embryos to achieve gene knockout. The technical procedures of microinjection followed Xu et al. 2025. The maintenance of injected embryos followed the procedure described by Neupane et al 2025.

### Genotyping and identification of mutants of target genes

For each female neonate hatched out of the injected embryos, we separated them and raised them to maturity. We call these G0 (generation 0) founders. The different broods of the founders are called brood 1, brood 2, and so on. Neonates from brood 1 or later broods are separated to establish a clonal culture for DNA extraction and genotyping. For the SCARLET gene in *D. magna*, we recorded the number of offspring with different eye pigment phenotypes (black vs. clear) for 5 broods. We genotyped 7-12 offspring from brood 1-3 each for all the G0 founders. For the other genes, we genotyped 1-5 offspring from brood 1 of the G0 founders.

To identify mutants from brood 1 or later brood individuals, we used an M13 fluorescence-tagged PCR approach for fragment analysis (Schuelke 2000). The PCR primers can be found in Supplementary Table 2. In each fragment analysis, we include the wildtype sample as a comparison. The detailed procedure of genotyping fragment analysis can be found in Neupane et al 2025.

## Results

### The phenotypic and genotypic diversity of SCARLET knockout mutants of D. magna

For the SCARLET gene, we obtained 5 G0 founders, 3 of which are a mosaic knockout phenotype (i.e., partial loss of black pigment) and 2 of which are clear-eyed (**Table 1**). For each founder, we examined the eye phenotype of the offspring from brood 1 to brood 4. Four of the founders produced 100% clear-eyed offspring in all 4 broods (**Table 1**), whereas one founder (Scar7) produced 1 (out of 8) and 2 (out of 27) clear-eyed offspring in broods 1 and 2, respectively, with brood 3 and 4 being all clear-eyed. This observation suggests that the primordial germ cells (PGCs) in all the founders were effectively edited by the RNP at the SCARLET locus. The number of unedited PGCs seemed to be limited in the founder Scar7 and were exhausted after two broods of reproduction.

**Table 1.**
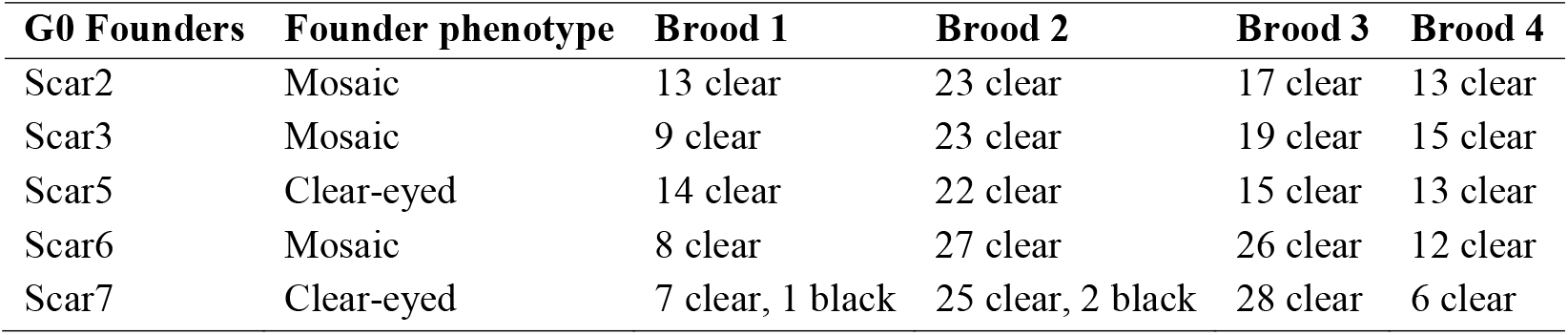
Summary of the number of black-eyed and clear-eyed neonates from SCARLET mutant founders. Only the phenotypes present in a brood are shown.

Notably, a diverse set of mutant genotypes is associated with the clear-eyed phenotype. In each brood of the G0 founders where 7-12 offspring were genotyped, all mutants showed biallelic genetic modifications, with the number of distinct genotypes at the scarlet locus ranged from 2 to 6 (**Figure 1**). Only one founder (Scar2) had the same two genotypes across 3 broods, suggesting that the RNP editing activity occurred in the 32-cell embryonic stage when 2 PGCs are present. All other founders gave rise to more than 2 distinct genotypes in their offspring, suggesting RNP editing activity did not occur in the one-cell embryonic stage and likely lasted until when more than 2 PGCs are present (later than the 32-cell embryonic stage).

**Figure 1.**
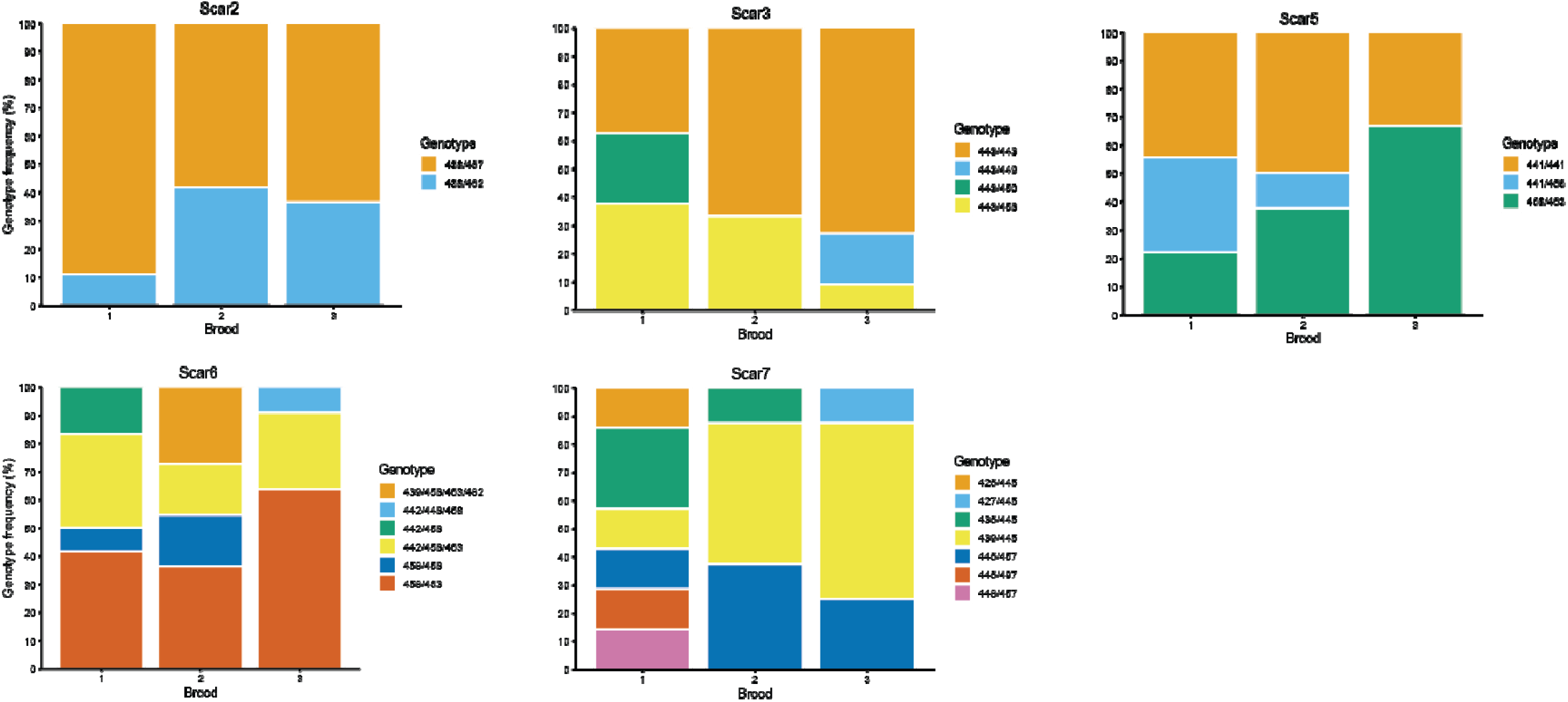
The summary of the percentages of different SCARLET mutant genotypes in the first three broods of 5 G0 founders. The wildtype genotype is 444/444.

Interestingly, two mutant genotypes with 3 and 4 alleles were detected in the offspring of the founder Scar6 (**Figure 1**), suggesting duplication likely occurred. These genotypes were verified through additional rounds of independent PCR and genotyping runs.

### *Genotypic diversity of knockout mutants in* D. magna

We performed gene knock-out experiments for 6 other genes in *D. magna* including DNMT3A, DNMT3B, CLOCK, PERIOD, PRDM8, DMRT1 (**Supplementary Table 1**). For DNMT3B, we identified 5 G0 founders and each one of them produced 1 unique biallelic mutant genotype in their brood 1 offspring where 3-5 offspring were genotyped (**Table 2**), suggesting efficient RNP editing activity at the 1-cell embryonic stage or before the first primordial germ cell was specified. For the other genes, we also found evidence that RNP editing activity took place in the 1-cell embryonic stage or before the specification of first primordial germ cell, with at least 1 founder identified for each gene producing a single unique genotype in their first brood (2-5 offspring genotyped). Interestingly, it is common to find multiple (2-5) different biallelic mutant genotypes in the first brood of the G0 founders, whereas wildtype genotype and monoallelic mutant genotypes were also present in CLOCK, PRDM8, and DMRT1 (**Table 2**). This set of observations suggest that the RNP editing was still active when multiple PGCs were present.

**Table 2.**
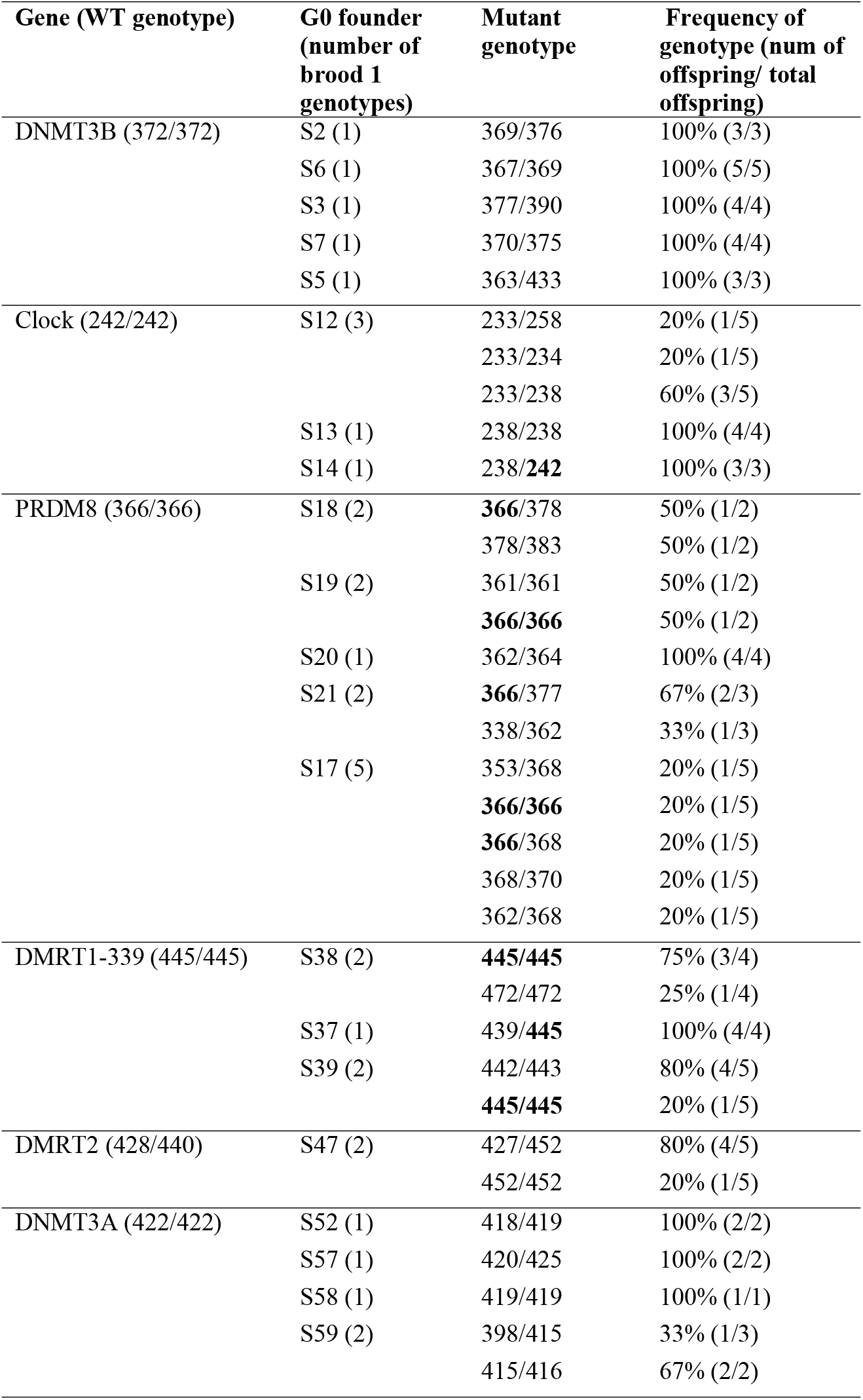
Summary of the identified genotypes in first brood of the founders from gene knockout experiments of six genes for *D. magna* LRV1 isolate. Bold numbers in the mutant genotypes represent wildtype alleles.

**Table 3.** Summary of the identified genotypes in first brood of the founders from gene knockout experiments of six genes for *D. pulex* Tex23 isolate. Bold numbers in the mutant genotypes represent wildtype alleles.

| Gene (WT genotype) | G0 founder<br>(number of brood<br>1 genotypes) | Mutant genotype | Frequency of genotype<br>(num of offspring/ total<br>offspring) |
| --- | --- | --- | --- |
| DFH (**400/400**) | X1 (2) | 393/394/395/396 | 75% (3/4) |
|  |  | 395/396/402/403 | 25% (1/4) |
|  | X2 (2) | 382/383/398/399 | 50% (1/2) |
|  |  | 382/383/388/394/399/**400** | 50% (1/2) |
|  | X12 (1) | 392/402 | 100% (2/2) |
|  | X16 (1) | 392/393 | 100% (2/2) |
|  | X18 (2) | 384/392 | 50% (1/2) |
|  |  | 393/394 | 50% (1/2) |
|  | X20 (1) | 388/399 | 100% (4/4) |
|  | X22 (1) | 398/399 | 100% (2/2) |
| Insulin-like (**470/470**) | X37 (1) | 461/461 | 100% (4/4) |

**Table 4.** Summary of the identified genotypes in first brood of the founders from gene knockout experiments of six genes for *D. sinensis* DJCL2 isolate. Bold numbers in the mutant genotypes represent wildtype alleles.

| Gene (WT genotype) | G0 founder (number of brood 1 genotypes) | Mutant genotype | Frequency of genotype (num of offspring/ total offspring) |
| --- | --- | --- | --- |
| ISM1 (**445/445**) | D9 (1) | 236/438 | 100% (4/4) |
| TM2D2 (**485/485**) | D43 (2) | 481/**485** | 67% (2/3) |
|  |  | **485/485** | 33% (1/3) |
|  | D41 (5) | 456/474 | 10% (1/10) |
|  |  | **485/485** | 20% (2/10) |
|  |  | 483/483 | 10% (1/10) |
|  |  | 481/**485** | 10% (1/10) |
|  |  | 453/**485** | 50% (5/10) |
|  | D50 (3) | 457/483 | 20% (1/5) |
|  |  | **485/485** | 60% (3/5) |
|  |  | 477/**485** | 20% (1/5) |
|  | D44 (2) | 477/487 | 67% (2/3) |
|  |  | 477/**485** | 33% (1/3) |
|  | IT2 (3) | **485/485** | 50% (2/4) |
|  |  | 480/**485** | 25% (1/4) |
|  |  | 346/**485** | 25% (1/4) |
|  | IT6 (1) | 387/427 | 100% (4/4) |
|  | IT5 (2) | **485/487** | 50% (1/2) |
|  |  | 483/487 | 50% (1/2) |
|  | IT7 (2) | **485/485** | 50% (1/2) |
|  |  | **485/500** | 50% (1/2) |

### *Genotypic diversity of knockout mutants in* D. pulex *and* D. sinensis

We knocked out the genes DFH and insulin-like receptor in the *D. pulex* isolate Tex23 (**Table 2, Supplementary Table 1**). The gene Insulin-like-receptor had the same genotype in all 4 offspring in brood 1. For DFH, we identified 4 G0 founders that produced one unique genotype in all offspring and 1 founder that gave rise to 2 different genotypes in the offspring. Similar to the observation of more than 2 alleles in the SCARLET mutants in *D. magna* LRV1, two founders gave rise to mutants with multiple mutant alleles that were verified through two rounds of independent PCR and genotyping.

For the *D. sinensis* isolate DJCL2, we knocked out two genes, ISM1 and TM2D2. For ISM1, we identified 1 founder that gave rise to a unique genotype in all of the 4 brood 1 offspring. For TM2D2, among the 8 founders that gave rise to mutant offspring, only 1 founder produced a unique biallelic mutant genotype in the brood 1 offspring, whereas the other founders produced multiple genotypes including biallelic and monoallelic mutant genotypes and the wildtype genotype in the brood 1 offspring.

These results are consistent with our observations in *D. magna*, suggesting that the CRISPR-Cas9 RNP activity in *Daphnia* embryos take places from as early as the 1-cell embryonic stage and more often when multiple PGCs are present.

## Discussion

Our prior work using CRISPR-Cas9 RNP to knock out the SCARLET gene in *D. pulex* (Xu et al. 2025) suggested that the RNP injected into asexual *Daphnia* embryos often leads to mosaicism in founder mutants and different mutant genotypes in their asexually produced offspring. It has not been clear whether this type of mosaicism was unique to the SCARLET gene or to the *D. pulex* species. To address this question and better understand the CRISPR-Cas9 RNP editing activity in the context of *Daphnia* asexual embryo development, this study presents CRISPR-Cas9 gene knockout experiments for 11 genes in three *D. pulex* species including *D. magna, D. pulex*, and *D. sinensis*. Based on extensive genotyping of the first brood of asexual offspring of founder mutants, this study reveals a few general patterns of RNP editing activity that are helpful for guiding future gene editing experiments in *Daphnia*.

First, mosaicism due to CRISPR-Cas9 gene editing is common for 9 out of the 11 surveyed genes in three *Daphnia* species, except for the genes DNMT3B and ISM1. Although oocytes are supposed to be genetically identical during the asexual reproduction of *Daphnia*, genetically distinct offspring from the first brood of the G0 founders (i.e., females hatched from the edited embryos) are present for most of the genes. This suggests that the G0 founders contain genetically distinct PGCs (primordial germ cells) due to the CRISPR-Cas9 RNP editing activity during embryo development.

Second, despite the efforts to deliver the CRISPR-Cas9 RNP during the 1-cell embryonic stage, in many cases RNP activity may only take place at later embryonic stages. Based on the identified number of genotypes in the brood 1 of G0 founders, we estimate that some of the gene editing activities affecting the germ cells occur at the 32-cell and later embryonic stage. Assuming the *Daphnia* embryo undergoes one round of cell division every 30 minutes (Sagawa et al. 2005), this means CRISPR-Cas9 RNP activity affecting the germ cells occurs at least 2.5 hours after injection.

Third, it is unclear why some genes such as DNMT3B and ISM1 did not show mosaicism, i.e., all the brood 1 offspring of G0 founders had the same genotypes, whereas other genes show extensive mosaicism. As our experiments were carried out in the same way, this indicates the RNP of these genes works more efficiently on the target sites at the 1-cell embryonic stage than that of the other genes, despite they were designed using the same rules and were in vitro tested for their DNA cutting activity prior to our gene editing experiments. It is likely other biological factors may be in play here, for example, the chromatin state of the CRISPR-Cas9 target sites. It has been demonstrated that a closed chromatin state can inhibit CRISPR-Cas9 editing in human cells (Daer et al. 2017). Thus, integrating chromatin state information into designing guide RNAs may yield a fruitful way of reducing mosaicism.

Furthermore, the genotyping data for the SCARLET gene for three consecutive broods (**Figure 1**) indicate that oocytes going into the same brood can originate from as many as 6 different lineages (e.g., brood 1 for Scar7). Also, we find evidence of turnover of oogonia cell lineages, with specific genotypes appearing in a subset of broods (e.g., Scar3, Scar5, Scar6, Scar7). The disappearance of early-brood genotypes (e.g., 441/458 genotype in broods1 and 2 for Scar5) in later broods suggests that exhaustion of oogonia of specific genotypic lineages.

We suggest that mosaicism resulted from CRISPR-Cas9 in *Daphnia*, although requiring rigorous genotyping to identify unique mutants, is a useful feature for generating multiple loss-of-function genotypes. Moreover, for CRISPR-Cas9-mediated gene knock-in experiments, which rely on homology dependent repair pathway and a homologous DNA template, mosaicism in germ cells may lead to multiple repair outcomes and enhance the chances of obtaining the correctly inserted DNA sequence at the target locus.

As this study relies on fragment length genotyping data to identify mutant genotypes, it is unclear whether the two guide RNAs for the same gene work equally efficiently in generating the mutant genotypes. It also remains unknown whether the two alleles of a gene are both mutated by CRISPR-Cas9 RNP. To gain insight into these aspects, we suggest an amplicon-sequencing approach for future studies. A sequencing approach could help resolve the likely duplication events in *D. magna* and *D. pulex* observed in the mutants of the SCARLET and DFH genes, which are not commonly considered as the undesirable outcomes of CRISPR-Cas9 genomic editing (Aussel et al. 2025).

## Acknowledgements

This work is supported by NSF EDGE grant 2220695/2324639, NIH grant R35GM133730, NSF CAREER grant MCB-2042490/2348390 to SX.

## Notes

### Competing Interest Statement

The authors have declared no competing interest.

