## Supplementary materials for "CRISPR-Cas9-induced genetic mosaicism in three species of the microcrustacean *Daphnia*"

**Supplementary Table 1.** Summary of target genes and target sites of guide RNAs. For *D. magna*, the gene IDs and locations refer to the *D. magna* genome assembly GCA_030254905.1(NCBI Accession: PRJNA777104). For *D. pulex*, the DFH gene refer to the genome assembly GCF_021134715.1 (NCBI Accession: PRJNA794129), whereas the insulin-like receptor refers to genome assembly GCA_911175335.1 (NACBI Accession: PRJEB46221).

| **Target Gene** | **Gene ID** | **sgRNAs** | **Genomic Location** | **Target Sequence (5´-3´)** |
| --- | --- | --- | --- | --- |
| ***D. magna*** |  |  |  |  |
| *SCARLET* | Dmagna032091 | *MasgRNA1* | Scaffold_8:2928389-2928412 (-) | GACGTCGTAACAGGCGTGGTGGG |
|  |  | *MasgRNA2* | Scaffold_8:2928625-2928647 (+) | CCTGGCGTGATTTGAGCGTCTAC |
| *CLOCK* | Dmagna027176 | clksgRNA3 | scaffold_7:7383606-7383625 (-) | GGTGGGCTATCAAGTTCAAA |
| *DNMT3A* | Dmagna014665 | *DNMT3A*-Mtase-sgRNA1 | scaffold_4:1379821-1379843 (-) | GAAGAGGTGTACGAACTCAA |
|  |  | *DNMT3A*-Mtase-sgRNA2 | scaffold_4:1379724-1379746 (+) | GGTCGTGGTACTCCACGACA |
| *DMRT2* | Dmagna027218 | DMRT2 | scaffold_7:7542111-7542130 (-) | ATGGACAGCGCTCTACGTTT |
|  |  | DMRT2.3 | scaffold_7:7541929-7541948 (-) | AAGAGTCATGGCTGCCCAGG |
| *DNMT3B* | Dmagna014617 | MA*DNMT3B*-sgRNA1 | scaffold_4:1221965-1221987 (-) | CCCCTCTTTTCTATCTAGCCAAA |
|  |  | MA*DNMT3B*-sgRNA2 | scaffold_4:1221777-1221799 (-) | CCATCCGTTTTTTAACGGTGGTC |
| *DMRT1* | Dmagna000339 | *DMRT1*dm | scaffold_1:1580546-1580565 (-) | TGTGCAAGAATCATGGCATC |
|  |  | *DMRT1*-2 | scaffold_1:1580730-1580749 (-) | TGGCCGAAATAGATGCCCTT |
| PRDM8 | Dmagna031900 | *prdm8sgRNA1* | scaffold_9:2135212-2135231 (-) | ATGGACACGACGATGGATAG |
|  |  | *prdm8sgRNA3* | scaffold_9:2135140-2135159 (-) | TTTGATGTGGCCTTTCTAAC |
| ***D. pulex*** |  |  |  |  |
| DFH | LOC124196108 | *sgRNA1* | NC_060017.1:531447-531466 (-) | CAGCGGAAAAAAGTTTAAAG |
|  |  | *sgRNA2* | NC_060017.1:531600-531619 (+) | GTCAACGATCGAGACCCATT |
| Insulin-like receptor | gene3866 | *sgRNA3.3* | CAJVNM010000001.1:5021727-5021747 | GCTGACAAACGTCAGTTATT |
|  |  | *sgRNA1.4* | CAJVNM010000001.1:5021659-5021679 | CGTCACTATTGTTCACAAGT |
| ***D. sinensis*** |  |  |  |  |
| ISM1 | Dsinen04828 | *sgRNA1* | chr3:1836045-1836064 (+) | GAGTGTGGGATCCTCTCAAC |
|  |  | *sgRNA3* | chr3:1836264-1836283 (-) | CAAAGAGAGACGGTATTTCT |
| TM2D2 | Dsinen04830 | *sgRNA2* | chr3:1842482-1842501 (+) | GTTGGAAAGCTCTTAACTCT |
|  |  | *sgRNA3* | chr3:1842443-1842462 (+) | GGCATTGATCGATTCTGTTT |

**Supplementary Table 2.** Summary of the PCR primers used for genotyping analysis. The forward primer contains M13 sequence at the 5´.

| **Target Gene** | **Primers** | **Primer Sequences (5´-3´)** |
| --- | --- | --- |
| *SCARLET* | MaseqF1-M13 | CACGACGTTGTAAAACGACGAGACCTTATAGGGCGGAAC |
|  | MaseqR | aacgtctagctgcaatacCATT |
| *DNMT3A* | MADNMT3A-F-M13 | CACGACGTTGTAAAACGACGTTCCGACTGGGCCCTTAAA |
|  | MADNMT3A-r | GATGACGTTGCGGATCGAAT |
| *DMRT2* | dmrt2fw1-M13 | *CACGACGTTGTAAAACGAC*ACAGCCAATCAGGACACGCAGC |
|  | dmrt2rv1(M13) | TCTTTGCCGCTCGACGACCAAC |
| *DNMT3B* | MADNMT3B-F | CACGACGTTGTAAAACGACCGCAGTTCAGTGAATCCGA |
|  | MADNMT3B-R | CAATTCTGTGCTTGTCCTATGG |
| *DMRT1* | DMRT1fw1-M13 | CACGACGTTGTAAAACGACTGTGCAGAATTCGGCCAGCGTT |
|  | DMRT1rv1(M13) | CTCTGTCTTGTTGCTGGGCCCG |
| PRDM8 | Prdm8_2fwd-M13 | CACGACGTTGTAAAACGACCAGATCGACCAACCAACTAACC |
|  | Prdm8_2-rv | GCGGCACATCTCAAATCTTCT |
| DFH | tex23dfhfwd1-M13 | CACGACGTTGTAAAACGACCGGGCAGCGTATAAATACCGATCGC |
|  | tex23dfhrv1 | TACGCGTGGGGCAGAATCGG |
| Insulin-like receptor | 3866-fw1-M13 | CACGACGTTGTAAAACGACTGGCCCAAGACACAGAATTTGACGA |
|  | 3866rv1(M13) | GCAGCAACATCAATTGTGGTCCACc |
| ISM1 | ism1-fw-M13 | CACGACGTTGTAAAACGACCCTCGGAGCGCAACGGAAGATT |
|  | ism1-rv(M13) | CTGGCCGATAGATGTCGACGCG |
| TM2D2 | SnTM2Fw2-M13 | CACGACGTTGTAAAACGACTTCATTTCAGGTACACAGGCCA |
|  | SnTm2rv2 | GAAACGGGAGCATGGGGGTTCG |
